# Comparing peptide identifications by FAIMS versus quadrupole gas-phase fractionation

**DOI:** 10.1101/2023.09.01.552989

**Authors:** Danielle A. Faivre, Christopher D. McGann, Gennifer E. Merrihew, Devin K. Schweppe, Michael J. MacCoss

## Abstract

High-field asymmetric waveform ion mobility spectrometry (FAIMS) coupled to liquid chromatography-mass spectrometry (LC-MS) has been shown to increase peptide and protein detections compared to LC-MS/MS alone. However, FAIMS has not been compared to other methods of gas-phase fractionation, such as quadrupole gas-phase fractionation, which could increase our understanding of the mechanisms of improvement. The goal of this work was to assess whether FAIMS improves peptide identifications because 1) gas-phase fractionation enables the analysis of less abundant signals by excluding more abundant precursors from filling the ion trap, 2) the use of FAIMS reduces co-isolation of peptides during the MS/MS process resulting in a reduction of chimeric spectra, or 3) a combination of both. To investigate these hypotheses, pooled human brain tissue samples were measured in triplicate using FAIMS gas-phase fractionation, quadrupole gas-phase fractionation, or no gas-phase fractionation on two Thermo Eclipse Tribrid Mass Spectrometers. On both instruments, our data confirmed prior observations that FAIMS increased the number of peptides identified. We further demonstrated that the main benefit of FAIMS is due to the reduced co-isolation of persistent peptide precursor ions, which results in a decrease in chimeric spectra.

## Introduction

Ion trap mass spectrometers have been the workhorse instruments in many protein mass spectrometry laboratories for years. What quadrupole ion trap mass spectrometers lack in resolution or mass accuracy, they make up in full scan sensitivity, MS/MS scan speed, and round-the-clock robustness. With the availability of orbitrap mass analyzers, we now have Fourier transform mass accuracy, resolving power, and dynamic range in benchtop capabilities and in combination with a quadrupole ion trap.^1,2^

An important strength of many ion trap mass spectrometers is the application of automatic gain control (AGC).^3^ AGC accumulates low abundance molecular species to fill the trap and adjusts the accumulation time based on the abundance of analyte ions. In an MS1 survey spectrum covering a wide *m/z* range, a single abundant peptide can compromise the dynamic range if the majority of the ions within the trap represent the abundant peptide. However, in an MS/MS acquisition, only molecular species within the isolated precursor *m/z* window are stored within the trap. This facilitates the accumulation of low abundance species in the presence of more abundant ions at different *m/z*. Thus, it is not uncommon to obtain high quality MS/MS spectra for peptides when there was no detectable precursor during the MS1 survey scan.^4^ Because the instrument dynamically adjusts the ion accumulation time to fill the trap, the MS/MS spectra can be of similar quality regardless of the abundance of the analyte in the mixture.

Because of AGC, the use of an ion filter to eliminate abundant peptides is particularly powerful when used in combination with an ion trap mass analyzer. Filtering high abundance molecular species from the analyte stream allows the mass spectrometer to accumulate and fill the trap with low abundance peptides. This idea was the basis of DREAMS reported by the Smith lab where a quadrupole was used to selectively exclude abundant *m/z* values from the ion cyclotron resonance cell to improve the dynamic range.^5^ Additionally, the Goodlett lab made use of gas-phase fractionation to isolate only a narrow *m/z* range in the acquisition of their MS1 survey scan.^6^ More recently, Meier *et al*. implemented BoxCar where the maximal orbitrap charge capacity could be accumulated over multiple narrow and discontinuous isolation windows^7^—the full mass range covered in subsequent scans. All of these methods enable the exclusion of abundant precursors and facilitate the use of longer injection times to improve the sensitivity of low abundant precursors.

High-field asymmetric waveform ion mobility spectrometry (FAIMS) can be used to decrease the complexity of the mixture before the ions enter the mass spectrometer. FAIMS was first used to measure peptides by Guevremont and coworkers^8–10^ and was later applied to a biological sample by Venne *et al*.^11^ The FAIMS interface separates ions by applying a high-voltage asymmetric waveform to a set of electrodes and allowing ions, entrained in a carrier gas, to flow through the gap between the electrodes. The ion separation occurs because ions have different mobilities in high and low electric fields. The asymmetry of the applied electric field, coupled with the field-dependent ion mobility, causes ions to acquire a net displacement perpendicular to their direction of motion and collide with one of the electrodes. A DC offset (compensation voltage or CV) can be applied to one electrode, which can counter this displacement and enable the transmission of ions.

The FAIMS device thus acts as a filter, transmitting only a portion of the total ions into the mass spectrometer—reducing the sample complexity, minimizing chemical noise, and increasing the dynamic range.^8–10^ Because FAIMS filters ions by differential mobility in a continuous fashion, low abundance ions can be accumulated using AGC in a similar way to using a quadrupole to isolate a subset of the *m/z* range. Our lab and others have previously demonstrated that the FAIMS CV can be stepped in a synchronized fashion with the acquisition of mass spectra, allowing ions at multiple selected differential ion mobilities to be measured in either separate or the same LC-MS runs.^11,12^ Recent improvements in FAIMS hardware have renewed interest in the technology.^13–15^ The hardware improvements have made FAIMS into a gas-phase fractionation method that can easily be performed prior to an ion trapping mass spectrometer.

While the speed which the FAIMS CV can be stepped is slower than stepping to different *m/z* regions with a quadrupole filter, an advantage of FAIMS is that the separation is at least partially orthogonal to *m/z*.^12^

While the improvement of FAIMS on the identification of peptides by data dependent acquisition is clear, the mechanism for how FAIMS improves the identifications is not well understood. Previous FAIMS experiments have benchmarked their experiments against analyses performed simply without FAIMS and never had an alternative gas-phase fractionation method as a control.^13^ Thus, it is not clear whether FAIMS acts just like other gas-phase fractionation methods by separating ions from the HPLC into separate bins—enabling the ion trap mass spectrometer’s use of AGC to fill longer and improve the sensitivity for lower abundant peptide precursors in the absence of abundant peptides. If FAIMS works simply by improving the dynamic range of the MS1 spectrum, then a gas-phase fractionation method that makes use of the quadrupole to isolate subsets of the *m/z* range should perform equally as well as, or better than, FAIMS. Thus, the goal of this work was to assess whether FAIMS improves peptide identifications because 1) gas-phase fractionation enables the analysis of less abundant signals by excluding more abundant precursors from filling the ion trap, 2) the use of FAIMS reduces co-isolation of peptides during the MS/MS process resulting in a reduction of chimeric spectra, or 3) a combination of both.

## Materials & Methods

### Mass Spectrometry

This experiment was designed to replicate a previous experiment,^13^ introduce a new control, and test the reproducibility of results on more than one instrument. Eluted peptides were analyzed with two Orbitrap Eclipse Tribrids. The two instruments examined different pooled human brain tissue samples with slightly different LC setups and gradients. Sample preparation has been described in detail previously,^16^ and the nanoLC conditions can be found in the Supporting Information. Experiments without FAIMS used a 240,000 resolving power MS1 survey scan, Standard AGC Target, and Auto Maximum Injection Time, followed by MS/MS of the most intense precursors for 1 second. The MS/MS analyses were performed by 0.7 *m/z* isolation with the quadrupole, normalized HCD (higher-energy collisional dissociation) energy of 30%, and analysis of fragment ions in the ion trap using the “Turbo” speed scanning from 200 to 1200 *m/z*. Dynamic exclusion was set to 10 seconds for the 1-hour analyses and was increased to 30 seconds for the 3-hour analyses. Monoisotopic precursor selection (MIPS) was set to Peptide, maximum injection time was 35 milliseconds, AGC target was 200%, unusual charge states (unknown, +1, or >+5) were excluded, the advanced peak determination was toggled on, and the internal mass calibration was off.

For FAIMS experiments, the settings were identical except the FAIMS device was used between the electrospray source and the mass spectrometer. FAIMS separations were performed with a 100 °C inner electrode temperature, 100 °C outer electrode temperature, 4.7 L/min FAIMS carrier gas flow, and −5000 V dispersion voltage (DV). The FAIMS carrier gas was N_2_ sourced from evaporated liquid nitrogen. For external stepping (i.e., single CV or single quadrupole fraction) experiments, the selected CV or quadrupole fraction was applied to all scans throughout the analysis. For internal stepping experiments, each of the 3 selected CVs or quadrupole fractions was applied to sequential survey scans. The MS/MS CV was always paired with the appropriate CV from the corresponding survey scan. For the 3-hour quantitative FAIMS experiments, the survey scan MS resolving power was reduced to 120,000 to permit a cycle time of 0.6 s. The 3 selected CVs (−50, -65, and -85) were chosen based on the results in Hebert *et al*.^13^ The 3 quadrupole fractions were sample-specific and were chosen based on splitting the number of peptide-like features in a normal LC-MS run into thirds.

### Data Analysis for Identifications

Raw files were converted into mzMLs using ProteoWizard’s msConvert.^17^ Peptides and proteins present in the samples were identified using Comet^18^ by searching against the human proteome plus common contaminants. The Comet search results were post-processed and a q-value was assigned to each PSM and each peptide using Percolator.^19^ The data was visualized using Limelight.^20^ All reported data are filtered at a false discovery rate (FDR) of 1%.

### mzML Splitting

The feature finding tools Hardklor^21^ and Bullseye^22^ need features to be present in multiple scans in a row to be considered persistent. To enable this analysis, we generated separate mzML files for each compensation voltage and quadrupole fraction. Available data conversion software such as msConvert^17^ can generate compatible mzML files, but it does not currently separate scans by different compensation voltages. Compatible mzML files were created from unseparated mzML files using a Python script developed in-house (https://github.com/uw-maccosslab/faims_vs_quadgpf). The script uses functionality provided via the pymzML module (https://github.com/pymzml/pymzML).^23^

### Data Sharing

All raw data, mzMLs, and other files used for analysis are available on Panorama Public (https://panoramaweb.org/faims_vs_quadgpf.url, ProteomeXchange ID: PXD043458). The database search results can be seen on Limelight (https://limelight.yeastrc.org/limelight/p/faims_vs_quadgpf). The figures can be reproduced using the open source code found on GitHub (https://github.com/uw-maccosslab/faims_vs_quadgpf).

## Results and Discussion

To improve the understanding of the mechanism of how FAIMS improves peptide identifications, we analyzed tryptic digests of human brain tissue (Supplementary Information).^16^ Samples were measured in triplicate using data dependent acquisition with FAIMS gas-phase fractionation (3 CV steps: -50, -65, -85 V), with quadrupole gas-phase fractionation (three selected ion monitoring *m/z* ranges), or without any gas-phase fractionation (Figure 1, right side). We also examined what we call external stepping versus internal stepping for the gas-phase fractionation methods (Figure 1, left side). For external stepping, a selected CV or limited mass range was applied to all scans throughout the analysis. For internal stepping experiments, each of the selected CVs or limited mass ranges was applied to sequential survey scans and MS/MS cycles. The MS/MS CV was always paired with the same CV from the corresponding MS1 survey scan.

**Figure 1.**
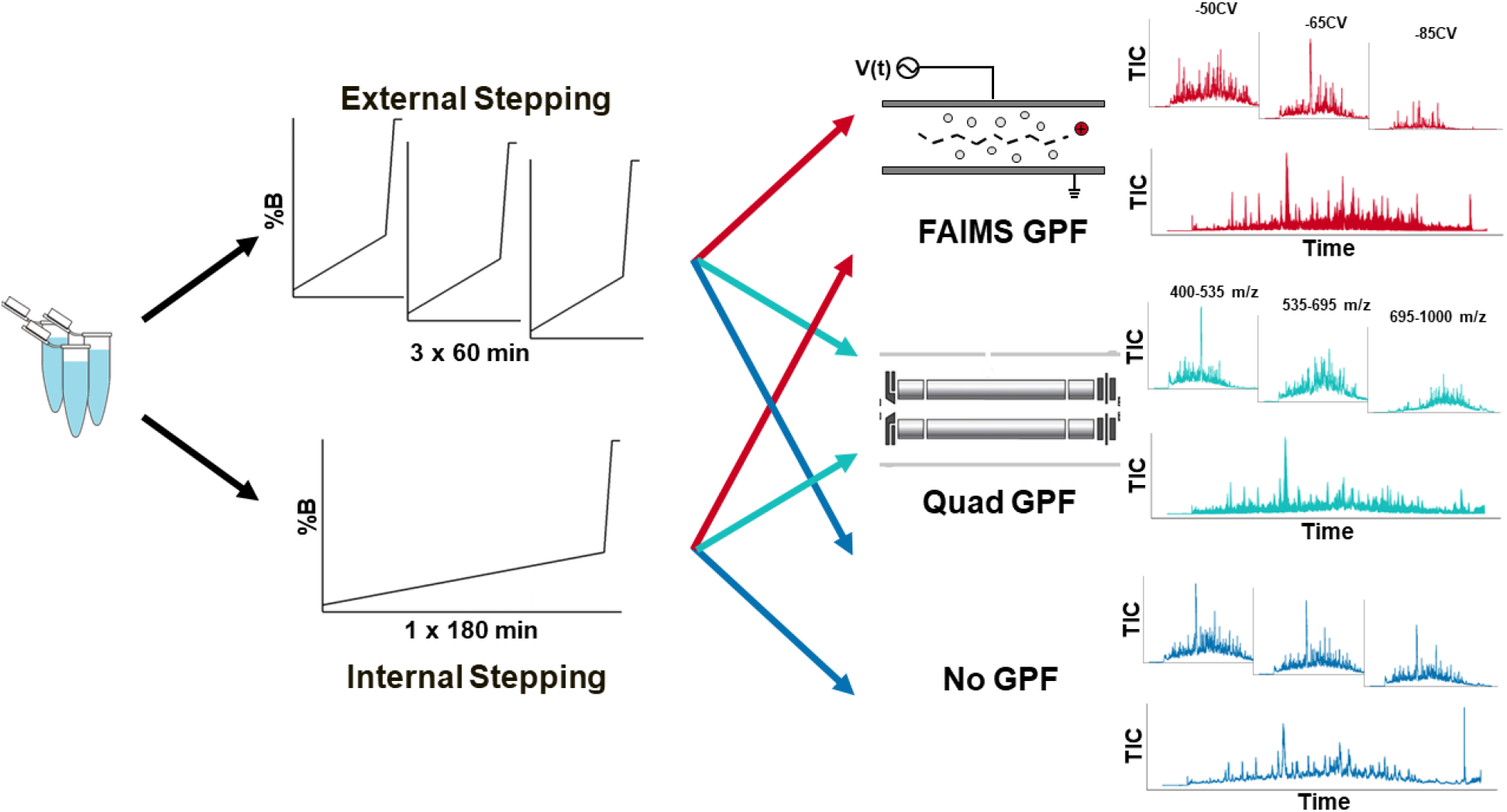
Scheme for the comparison of different gas-phase fractionation (GPF) methods to improve data dependent acquisition mass spectrometry. Pooled human brain tissue samples were measured in triplicate using external stepping (three 1-hour runs) and internal stepping (one 3-hour run). The experiments used FAIMS gas-phase fractionation (3 CV steps: -50, -65, -85 V), quadrupole gas-phase fractionation (3 SIM quad regions), or no gas-phase fractionation (right side). Internal stepping applied each of the 3 selected CVs or quadrupole fractions to sequential survey scans in a run while external stepping applied the selected CV or quadrupole fraction to all scans in a run.

Previously, the internal stepping experiments outperformed the external stepping experiments for peptide and protein detections^13^. This trend was seen in the dataset collected with Instrument 1, but it was not statistically significant in all cases (Figure 2a). This result was expected because the instrument can quickly switch between different CVs internally while the external stepping experiments are limited by the dead column loading time at the beginning of each run and the column re-equilibration time at the end of the gradient—each injection of a sample consumes more material and adds significant time overhead. For Instrument 2, this trend was not seen (Figure 2b). This difference may be the result of the optimized LC gradient reducing dead time at the beginning of each run (Figure S1). While internal stepping may not have a significant improvement in identifications, the method is better than external stepping because the multiple FAIMS gas-phase fractionation performed significantly better (q-value < 0.05) in peptide and protein detections than quadrupole gas-phase fractionation or the absence of gas-phase fractionation on both instruments (Figure 2). While the number of detections was similar within the triplicate of a method, there was poor overlap of identifications within the triplicate of a method and between the three methods due to the stochasticity of data dependent acquisition (Figures S2 & S3).

**Figure 2.**
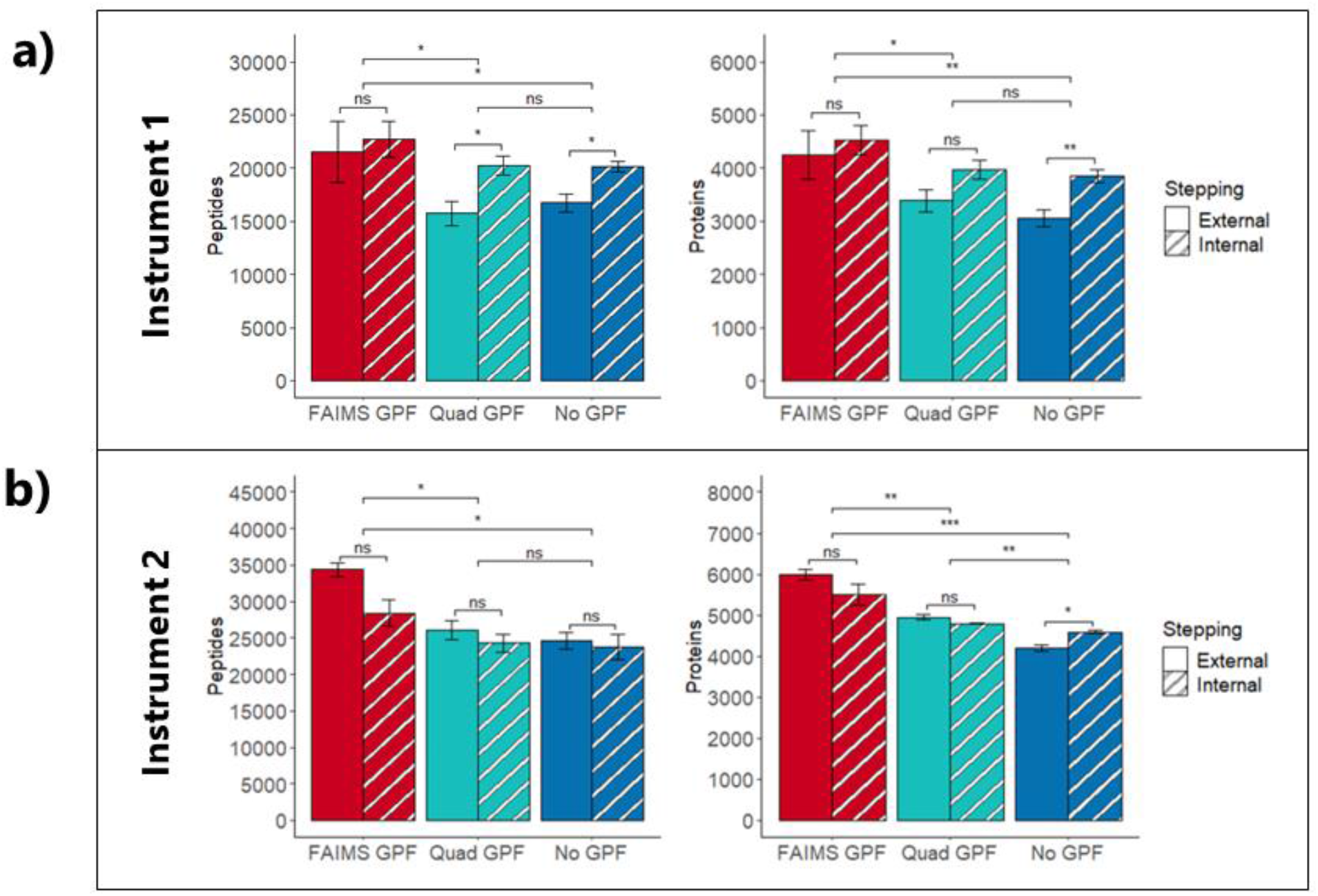
Peptide and protein counts for internal and external stepping gas-phase fractionation experiments. On both instruments, FAIMS gas-phase fractionation (red) performed significantly better (q-value < 0.05) in peptide (left) and protein (right) detections than quadrupole gas-phase fractionation (teal) or the absence of gas-phase fractionation (blue). The internal (striped) and external (solid) stepping results were not consistent between both runs needed for external stepping require more sample and additional time for gradient preparation, column equilibration, and sample loading.

We questioned if the lower number of identifications by quadrupole gas-phase fractionation was due to an underfilling of the ion trap, so we examined the ions measured in each method’s MS2 spectra. We observed that there was a larger difference in ion filling between FAIMS and non-FAIMS LC-MS (Figure 3a) than observed previously by Hebert *et al* (Figure S4).^13^ To confirm that the results from our initial experiment on an Orbitrap Eclipse was not an artifact of a single instrument, we repeated the experiments on a second Orbitrap Eclipse. The data from Instrument 2 (Figure 3b) minimized these concerns because the data between the two tribrids were similar.

**Figure 3.**
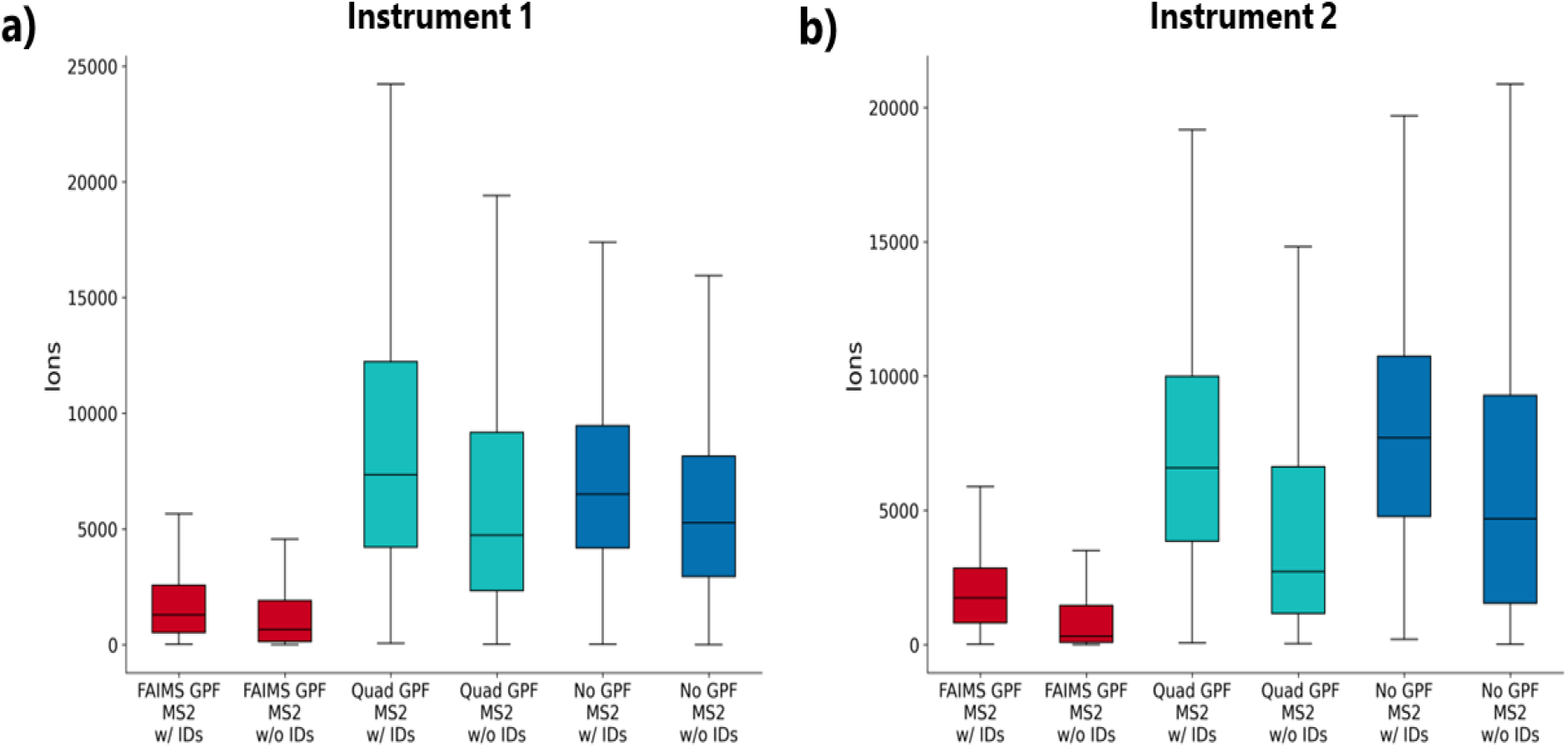
The number of ions measured in MS2 spectra for the respective LC-MS/MS runs. The number of ions were estimated by taking the total signal in each spectrum and multiplying by the ion injection time using the msions Python package. Boxplots demonstrate the median number of ions (middle line), extend from the first quartile to the third quartile (Q3 - Q1 = Interquartile Range or IQR), and have whiskers that extend to the minimum/maximum or 1.5 x IQR, whichever is less. The data show that FAIMS (red) underfills the ion trap more often than quadrupole gas-phase fractionation (teal) or normal LC-MS (blue). On average, the MS2 spectra with a larger number of ions are assigned a peptide identification with a low q-value at a higher frequency than spectra with fewer ions.

To assess if gas-phase fractionation enables the analysis of less abundant signals by excluding more abundant precursors from filling the ion trap, we examined the number of persistent peptide-like features found with each method. An advantage of quadrupole gas-phase fractionation is that, unlike FAIMS gas-phase fractionation, the MS1 features are distinct between fractions (Figure S5). This limited redundancy should increase the likelihood of selecting a low-intensity precursor for fragmentation.

The data collected using FAIMS had a larger percentage of low intensity MS1 precursors identified by MS/MS (Figure 4a), but the intensities of the features were also significantly lower (Figure 4b). This agreed with the underfilling of the ion trap shown in Figure 3. To determine how individual features were affected by the 3 methods, we examined the features that were identified by all 3 methods in one instrument batch. Compared to no gas-phase fractionation, FAIMS gas-phase fractionation lowered the intensity of over 90% of the shared identified features (Figure 4c) while quadrupole gas-phase fractionation increased the intensity of 90% of them (Figure 4d). These results support hypothesis 1 for quadrupole gas-phase fractionation, but not for FAIMS gas-phase fractionation.

**Figure 4.**
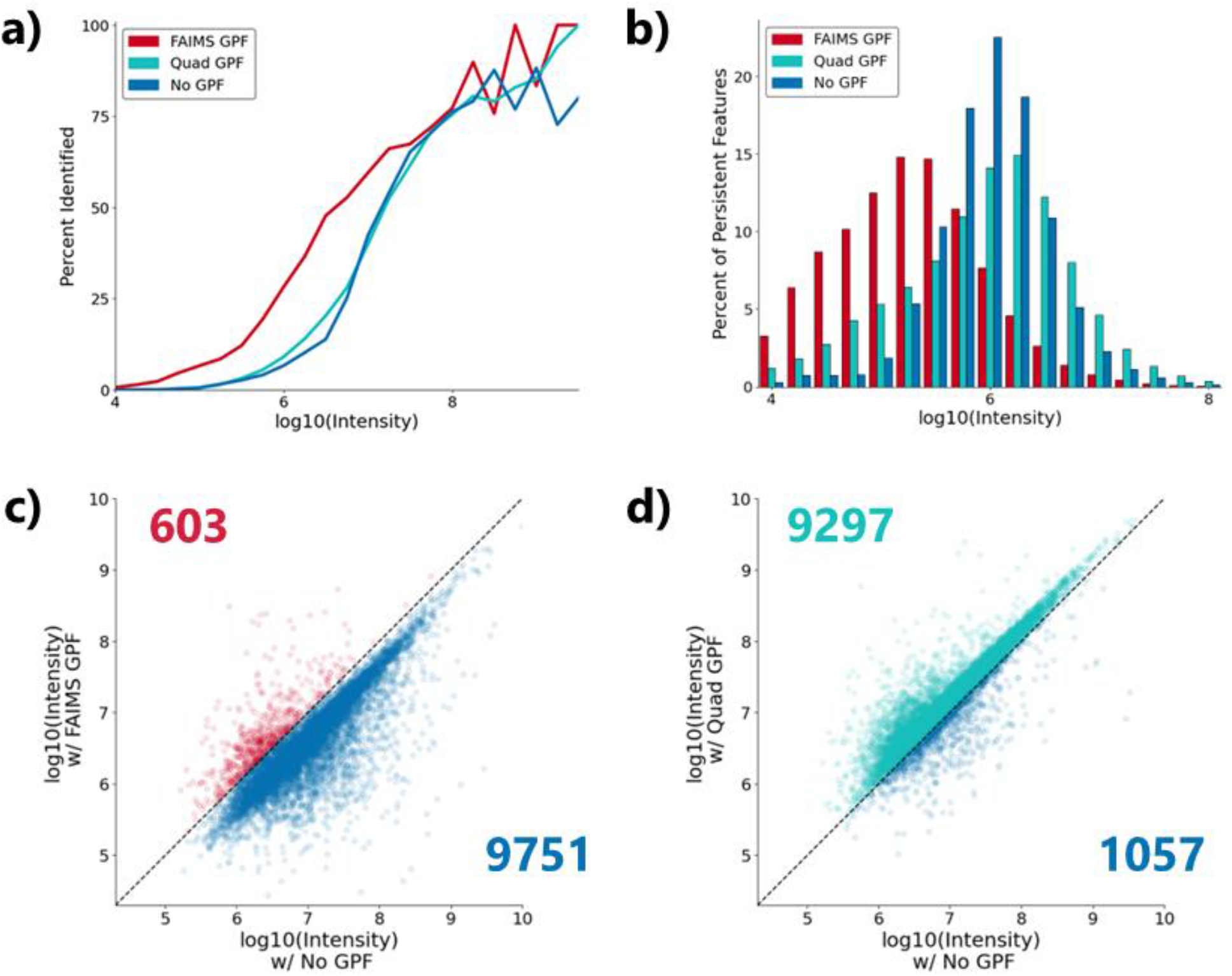
Comparison of persistent peptide-like features found with different gas-phase fractionation methods. Persistent peptide-like MS1 features were determined using Hardklor and Kronik. a. The percent of features identified based on their apex intensities (log10). b. The density of features in each intensity bin (log10). c. Comparison of the apex intensities of the same identified peptide features in a FAIMS gas-phase fractionated experiment (red) and experiment without gas-phase fractionation (blue). d. Comparison of the apex intensities of the same identified peptide features in a quadrupole gas fractionated experiment (teal) and experiment without gas-phase fractionation (blue).

To explore if the use of FAIMS reduces co-isolation of peptides during MS/MS and results in a reduction of chimeric spectra, we examined the number of persistent precursors present within quadrupole isolation windows. An isolation window is the narrow *m/z* range that is analyzed by the quadrupole during an MS2 spectrum acquisition. Non-chimeric spectra would only have one precursor in each MS2 spectrum and have a better chance of being identified because the identification rate of chimeric spectra can be 2-fold lower than non-chimeric spectra.^24^ Based on this criterion, FAIMS gas-phase fractionation successfully reduced chimeric spectra compared to quadrupole gas-phase fractionation and no gas-phase fractionation (Figure 5a & b). Quadrupole gas-phase fractionation also had fewer chimeric spectra than no gas-phase fractionation, but the improvement was smaller than with FAIMS. To investigate how FAIMS affects the chimeric spectra, we checked the relative intensities of the precursors in the isolation windows. On both instruments, the most intense precursor within an isolation window has a higher relative intensity for a higher percentage of FAIMS MS2 spectra (Figure 5c & d). This supports FAIMS improving the chances of identifying spectra that are chimeric in addition to reducing the number of chimeric spectra. We believe the different distributions seen between Instrument 1 and 2 could be explained by the sample and LC gradients being different. The data collected with Instrument 2 appeared to separate the peptides better across the elution time (Figure S1).

**Figure 5.**
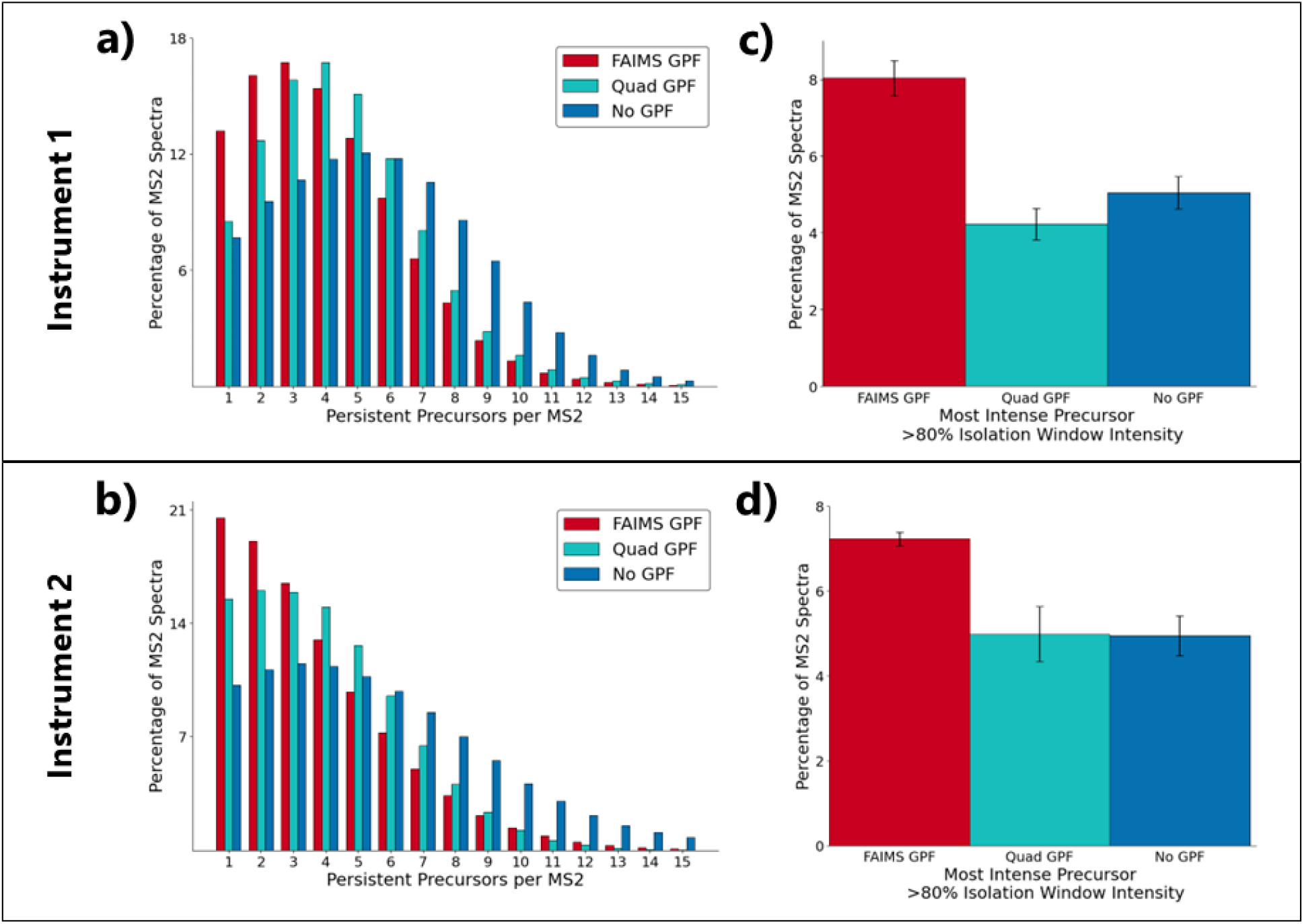
Investigating chimeric spectra with different gas-phase fractionation methods. a. Percentage of MS2 spectra with multiple precursors for Instrument 1. b. Percentage of MS2 spectra with multiple precursors for Instrument 2. c. Percentage of MS2 spectra where the most intense precursor had >80% of the summed isolation window intensity for Instrument 1. d. Percentage of MS2 spectra where the most intense precursor had >80% of the summed isolation window intensity for Instrument 2. FAIMS gas-phase fractionation reduced the number of chimeric spectra compared to quadrupole gas-phase fractionation and no gas-phase fractionation and improved the relative intensity of the most intense precursor in spectra that were chimeric.

## Conclusions

Overall, the results supported the use of FAIMS gas-phase fractionation over either quadrupole gas-phase fractionation or no gas-phase fractionation for this type of experiment. FAIMS reduced the co-isolation of persistent precursors during the MS/MS process, which resulted in a reduction of chimeric spectra, even though FAIMS gas-phase fractionation was less efficient at transmitting ions than quadrupole gas-phase fractionation. The lower ion transmission led to FAIMS lowering the intensity of over 90% of the shared identified features. We confirmed these results on two Thermo Eclipse Tribrid Mass Spectrometers while using two separate FAIMS interfaces, slightly different chromatography gradients, similar samples, and internal and external stepping. While internal stepping may not have a statistically significant improvement in identifications, the method should be chosen because the multiple runs needed for external stepping require more sample and additional time for each gradient preparation, column equilibration, and sample loading.

## Acknowledgements

This work was supported in part by grants RF1 AG053959, U19 AG065156, and R24GM141156 from the National Institutes of Health. DAF was supported by grant F31 AG066318. We would also like to thank Matthew Bush for providing feedback, and we appreciate Brian Connolly and Mike Riffle’s help with Cromwell and Limelight, respectively.

