## Supplementary Information for "Comparing peptide identifications by FAIMS versus quadrupole gas-phase fractionation"

### SUPPLEMENTARY FIGURES

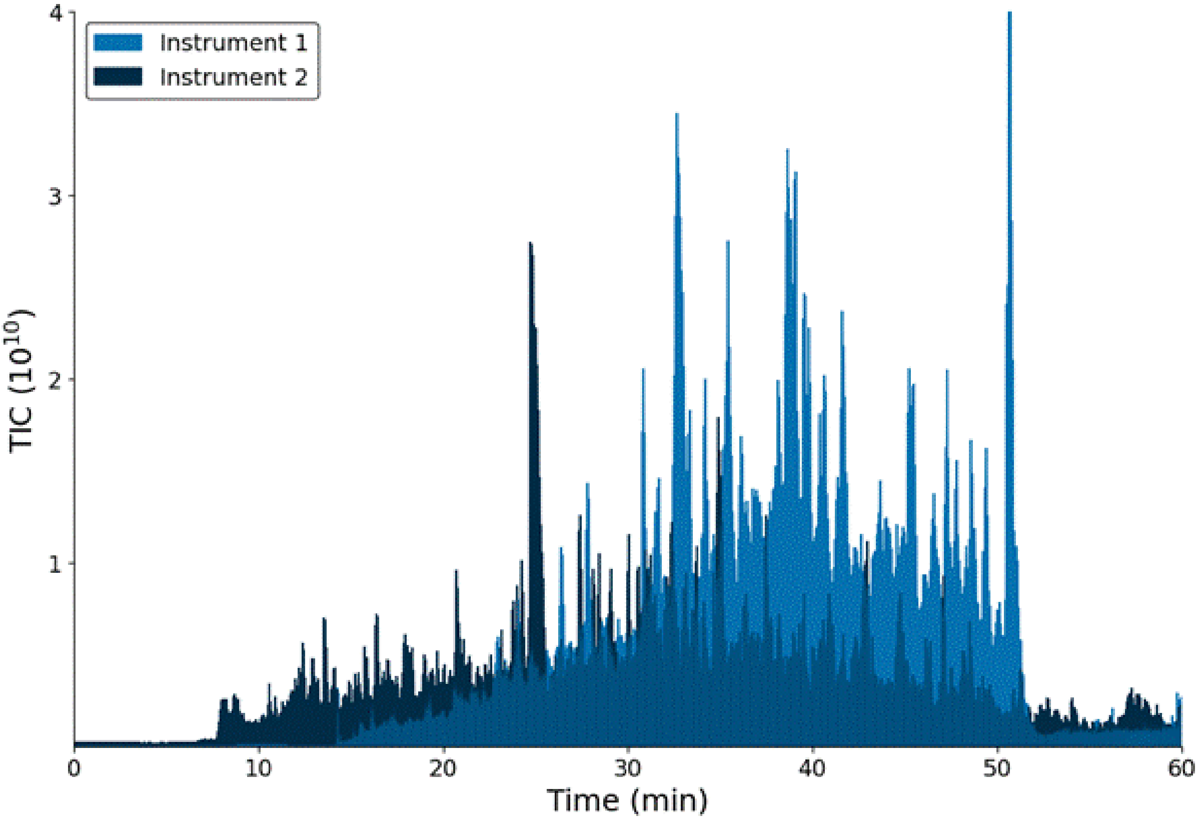


**Figure S1.** Example TICs for gradients used with Instrument 1 and Instrument 2.

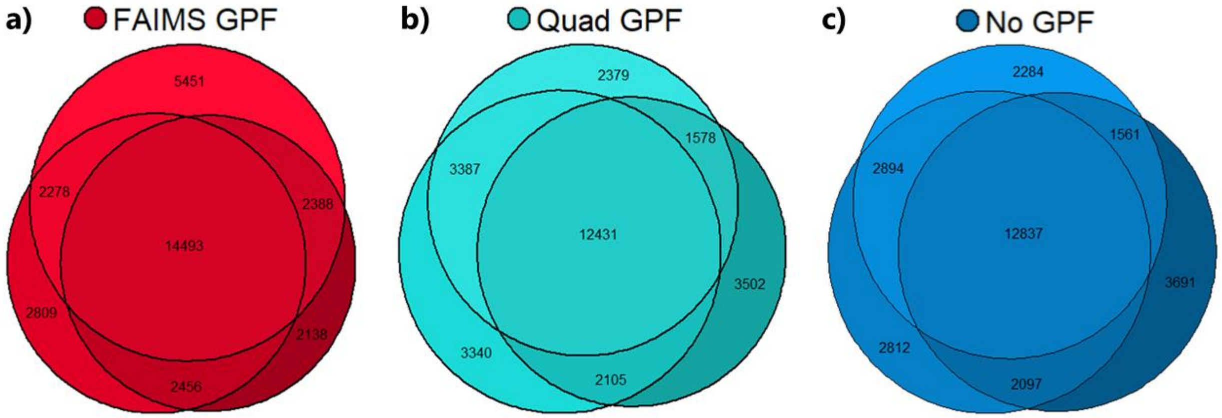


**Figure S2.** Overlap of identified peptides between the triplicate data dependent acquisition runs with FAIMS gas-phase fractionation (internal stepping), quadrupole gas-phase fractionation (internal stepping), and no gas-phase fractionation.

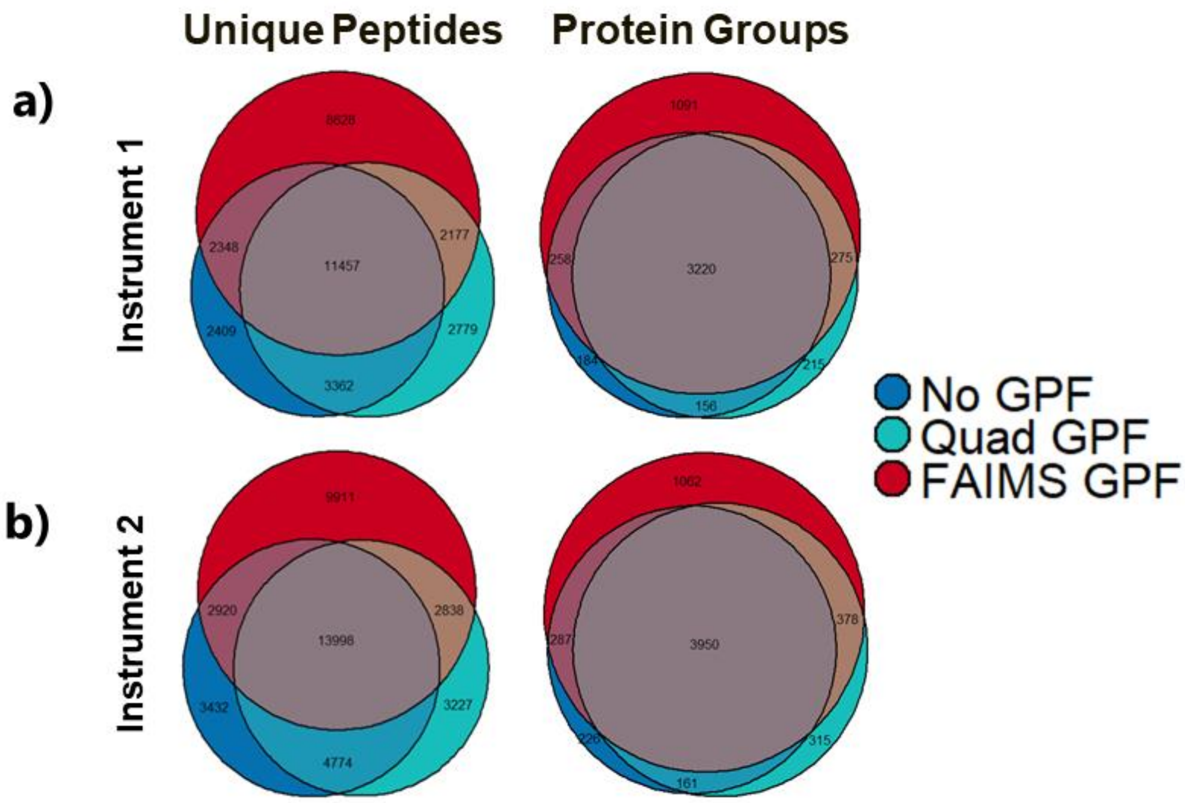


**Figure S3.** Overlap of identified peptides (left) and proteins (right) between FAIMS gas-phase fractionation, quadrupole gas-phase fractionation, or no gas-phase fractionation.

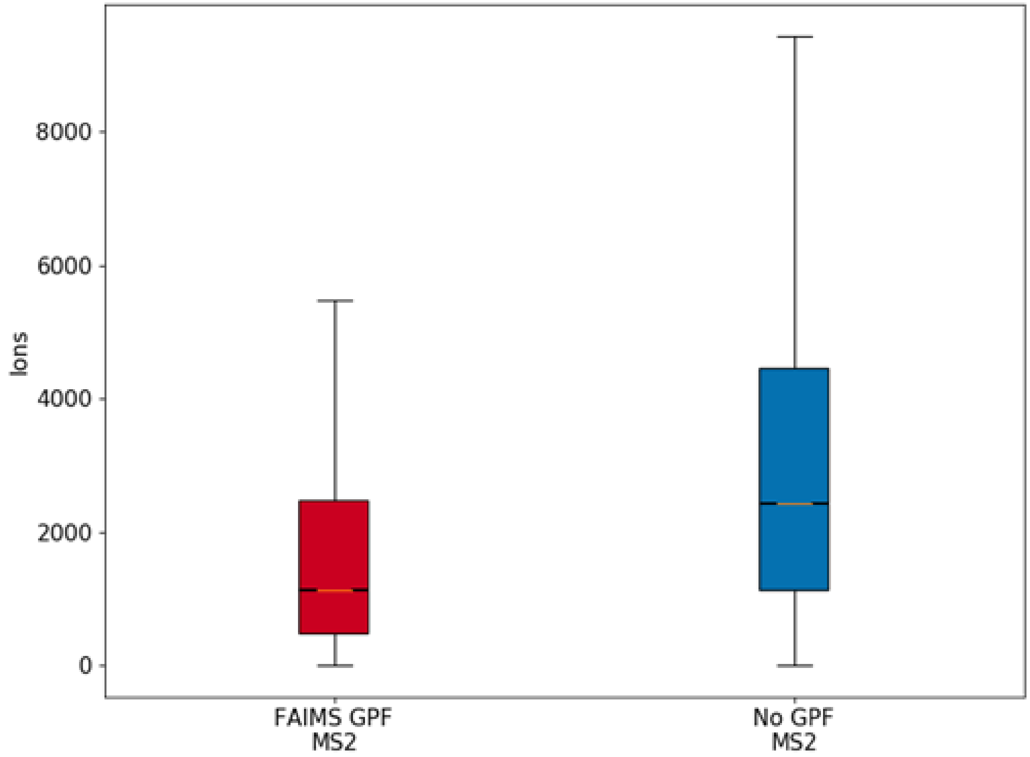


**Figure S4.** Ions injected for each MS2 spectra in Hebert *et al*.

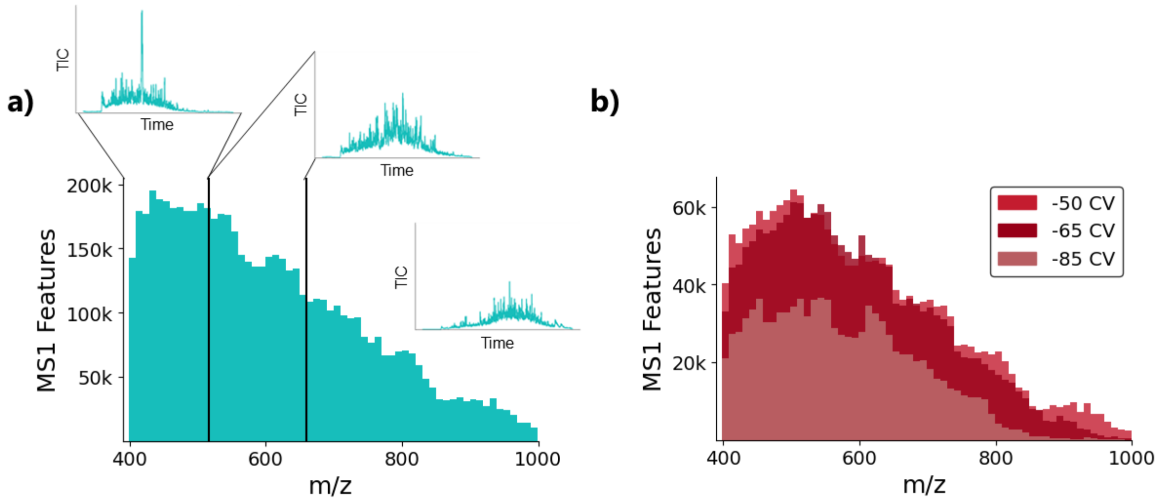


**Figure S5.** Overlap in *m/z* space for quadrupole (a) and FAIMS (b) gas-phase fractionation.

### SUPPORTING INFORMATION

#### Tryptic digestions.

Briefly, two 25 µm frozen sections of human brain tissue were resuspended in 120 µl of lysis buffer containing 5% SDS, 50mM triethylammonium bicarbonate (TEAB), 2mM $MgCl_2$, 1X HALT phosphatase, and protease inhibitors. The suspension was vortexed and briefly sonicated with a Fisher sonic dismembrator model 100 set to setting 3 for 10 s. A microtube was loaded with 30 µl of lysate and capped with a micropestle. The sample was homogenized with a Barocycler 2320EXT (Pressure Biosciences Inc.) for a total of 20 minutes at 35°C with 30 cycles of 20 seconds at 45,000 psi and 10 seconds at atmospheric pressure.

Protein concentration of the homogenate was measured with a BCA assay. Fifty micrograms were added to a process control of 800 ng of yeast enolase protein (Sigma), reduced with 20 mM DTT, and alkylated with 40 mM IAA. The lysate was prepared for S-trap column (Protifi) cleaning by adding 1.2% phosphoric acid and 350 µL of binding buffer (90% Methanol, 100 mM TEAB). The acidified lysate was bound to column incrementally, followed by 3 wash steps with binding buffer to remove SDS, 3 wash steps with 50:50 methanol:chloroform to remove lipids, and a final wash step with binding buffer. Trypsin (1:10) in 50mM TEAB was added to the S-trap column for digestion at 47°C for one hour. Hydrophilic peptides were eluted with 50 mM TEAB and hydrophobic peptides were eluted with a solution of 50% acetonitrile in 0.2% formic acid. Elutions were pooled, speed vacuumed and resuspended in 0.1% formic acid. For more sample details, see Merrihew *et al.*[16]

**NanoLC conditions.**

On both instruments, 1 µg of sample was loaded into the system. For instrument 1, a Thermo Easy-nLC 1200 was used with a 30 cm fused silica pulled tip column (New Objective, 75 µm inner diameter) and a 4 cm fused silica (150 µm inner diameter) Kasil1 (PQ Corporation) frit trap. The trap and column were loaded with 3 µm Reprosil-Pur C18 (Dr. Maisch) reverse-phase resin. Buffer A was 0.1% formic acid in water and buffer B was 0.1% formic acid in 80% acetonitrile. The 60-minute LC gradient was 2 to 7% B in 1 minute, 7 to 40% B over 40 minutes, 40 to 60% B over 5 minutes, 60 to 98% B over 5 minutes, a 5 minute wash at 98% B, a return to 2% B in 1 minute, and a 3 minute equilibration at 2% B. The 180-minute LC gradient was 2 to 7% B in 1 minute, 7 to 40% B over 160 minutes, 40 to 60% B over 5 minutes, 60 to 98% B over 5 minutes, a 5 minute wash at 98% B, a return to 2% B in 1 minute, and a 3 minute equilibration at 2% B. Peptides were eluted from the column with a 50°C heated source (CorSolutions) and electrosprayed into a Thermo Eclipse Tribrid Mass Spectrometer with the application of a distal 3 kV spray voltage. Application of the mass spectrometer and LC solvent gradients were controlled by ThermoFisher Xcalibur (version 3.3).

For instrument 2, 1 µg of sample was loaded into the system on an in-house pulled 30 cm C18 (Thermo Accucore, 2.6 Å, 150 µm) column. Buffer A was 5% acetonitrile/0.125% formic acid and buffer B was 0.125% formic acid in 95% acetonitrile. The 60-minute LC gradient was 4 to 35% B over 55 minutes and 35 to 100% B in 5 minutes. The 180-minute LC gradient was 4 to 35% B over 175 minutes and 35 to 100% B in 5 minutes. Both gradients had a 5 minute wash at 100% B. Peptides were eluted from the column and electrosprayed into a Thermo Eclipse Tribrid Mass Spectrometer with the application of a distal 3 kV spray voltage. Application of the mass spectrometer and LC solvent gradients were controlled by ThermoFisher Xcalibur (version 3.5).
